# The impact of α-synuclein overexpression on nigrostriatal transmission

**DOI:** 10.64898/2026.09.15.751842

**Authors:** Nikolas Bergum, Andrew G. Yee, Robert H. Edwards, Christopher P. Ford

## Abstract

Accumulation of wild-type α-synuclein in dopamine neurons of the Substantia Nigra *pars compacta* (SNc) is a pathological hallmark of Parkinson’s disease (PD), which commonly precedes neurodegeneration. Yet how α-synuclein accumulation affects nigrostriatal neurotransmission early in the progression of PD remains poorly understood. In addition to axonal dopamine release, SNc neurons also co-release GABA and glutamate from striatal axon terminals and release dopamine from their somatodendritic region. However, the selective vulnerability of these distinct functions to increased α-synuclein remains largely unexplored. Here, we selectively overexpressed wild-type human α-synuclein in SNc dopamine neurons to examine impacts on nigrostriatal and somatodendritic transmission and correlated these changes to impairments in locomotion. We found that dopamine release in the dorsolateral striatum was robustly reduced, while transmission to postsynaptic medium spiny neurons remained intact. Glutamate co-release from dopamine terminals onto both medium spiny neurons and cholinergic interneurons was selectively impaired, while GABA co-release was unaffected. Conversely, somatodendritic dopamine release within the SNc was completely abolished by α-synuclein overexpression. Together, these findings identify selective deficits in nigrostriatal neurotransmission associated with wild-type α-synuclein accumulation prior to overt degeneration and raise the possibility of compensatory adaptations shaping the early functional consequences of α-synuclein pathology in PD.

## Introduction

Parkinson’s disease (PD) is a neurodegenerative movement disorder with the pathological hallmarks of progressive degeneration of dopamine neurons in the substantia nigra pars compacta (SNc) and the formation of Lewy bodies within surviving neurons (Surmeier et al., 2017; Sharma and Burré, 2023). Lewy bodies are intraneuronal inclusions whose major component is the presynaptic protein α-synuclein (Spillantini et al., 1997). Rare familial forms of PD arise from point mutations or gene multiplications in *SNCA*, the gene encoding α-synuclein, indicating that excess or aberrant α-synuclein can be sufficient to cause disease (Polymeropoulos et al., 1997; Krüger et al., 1998; Singleton et al., 2003; Chartier-Harlin et al., 2004; Zarranz et al., 2004). However, the vast majority of PD cases are idiopathic where wild-type α-synuclein, rather than mutated forms, are elevated and accumulate (Sulzer and Edwards, 2019; Sharma and Burré, 2023). How wild-type α-synuclein accumulation disrupts neuronal function prior to overt degeneration however remains poorly understood.

α-synuclein is enriched at presynaptic terminals, where it associates with synaptic vesicle membranes and has been linked to the regulation of neurotransmitter release and SNARE complex assembly (Sulzer and Edwards, 2019; Sharma and Burré, 2023). Despite this, the function of α-synuclein at the synapse remains poorly understood. Overexpression of α-synuclein has been shown to impair neurotransmitter release across multiple systems, and α-synuclein-induced dopamine release deficits can emerge in the absence of SNc cell loss, suggesting that synaptic dysfunction precedes neurodegeneration (Nemani et al., 2010; Janezic et al., 2013; Parra-Rivas et al., 2023). However, because many experimental models rely on mutated forms of α-synuclein or induce overt degeneration, the early synaptic consequences of wild-type α-synuclein accumulation, as occurs in the majority of PD patients, have been difficult to isolate. To address this, we previously developed a cell-type-specific viral approach to overexpress wild-type human α-synuclein (hαSyn) selectively in midbrain dopamine neurons, producing progressive pathology and motor deficits in the absence of SNc cell loss (Barcomb et al., 2025). Here we used this model to determine how wild-type α-synuclein accumulation selectively alters the distinct neurotransmitter systems of nigrostriatal dopamine neurons.

SNc dopamine neurons are the principal source of striatal dopamine, which is critical for the control of voluntary movement. However, in addition, dopamine axons in the dorsal striatum co-release multiple transmitters including GABA and glutamate from a subset of terminals (Sulzer et al., 1998; Dal Bo et al., 2004; Hnasko et al., 2010; Tritsch et al., 2012; Kim et al., 2015; Zych and Ford, 2022; Chuhma et al., 2023). In a model of PD based on mutant A53T α-synuclein, deficits in GABA co-transmission at dopamine synapses were found to precede reductions in dopamine release itself, suggesting that co-release substrates may be differentially sensitive to α-synuclein-related pathology (Kim et al., 2023). In addition to striatal axonal release, SNc dopamine neurons release dopamine from their somatodendritic compartment, which activates D2 autoreceptors and has been proposed to play an important role in regulating motor output (Björklund and Lindvall, 1975; Geffen et al., 1976; Beckstead et al., 2004) and loss of somatodendritic dopamine release in SNc neurons may contribute to early parkinsonian motor deficits (González-Rodríguez et al., 2021; Muñoz et al., 2025). Whether wild-type α-synuclein overexpression differentially impairs GABA, glutamate, and dopamine neurotransmission from SNc dopamine neurons has not been examined.

Here, using ex vivo electrophysiology alongside characterization of motor impairment we investigated how α-synuclein accumulation affects each of these neurotransmitter systems. As α-synuclein overexpression produced motor impairments, we examined whether any deficits in nigrostriatal transmission correlated with these behavioral changes. We found that α-synuclein overexpression produced a selective deficit in bulk dopamine release and glutamate co-release in the striatum, as well as a near complete loss of somatodendritic dopamine transmission within the SNc. Together, these findings provide new insight into the early functional consequences of α-synuclein accumulation in dopamine neurons, and potentially into what drives early motor dysfunction in PD.

## Methods

### Animals

All procedures were approved by and performed in accordance with guidelines of the Institutional Animal Care and Use Committee (IACUC) at University of Colorado School of Medicine. Animals were group-housed in a temperature- and humidity-controlled environment on a 12 h/12 h light-dark cycle, with water and food available ad libitum, and experiments were conducted during the light phase. Both male and female 3-6 month-old DAT-IRES-Cre heterozygote mice (IMSR_JAX: 006660, Slc6a3IRES-Cre) were used in experiments. Ai32 mice (IMSR_JAX: 012569, RCL-ChR2(H134R)/EYFP) were crossed with DAT-IRES-Cre mice to express ChR2 in dopamine cells.

### Stereotaxic injection

For stereotaxic viral injections, postnatal 2-4 month-old male and female DAT-IRES-Cre; Ai32 heterozygote mice (DAT-Cre^(+/-)^, Ai32^(+/-)^) or DAT-Cre (DAT-Cre^(+/-)^) heterozygote mice were anesthetized with isoflurane and mounted in a stereotaxic frame (Kopf Instruments). For histology experiments, a Nanoject iii (Drummond Scientific) was used to inject 300 nL of AAV5.EF1a.DIO.IRES.mCh (AAV.mCh) into SNc of one hemisphere, while 300 nL of AAV5.EF1a.DIO.wthαSyn.IRES.mCh (AAV.hαSyn) was injected into the opposite hemisphere of DAT-Cre^(+/-)^ or DAT-Cre^(+/-)^, Ai32^(+/-)^ mice. For all voltammetry and behavioral experiments, 300 nL of AAV.mCh or 300 nL of AAV.hαSyn were bilaterally injected into the SNc of DAT-Cre^(+/-)^, Ai32^(+/-)^ mice. For the slice electrophysiology, DAT-Cre^(+/-)^, Ai32^(+/-)^ mice were injected with 300 nL of AAV.mCh or 300 nL of AAV.hαSyn into the SNc as well as 500 nL (1:4 dilution) of AAV9.hSyn.tdTomato.T2A.mGIRK2-1-A22A.WPRE.bGH (AAV.GIRK2) into the dorsal striatum of the same hemisphere. The following coordinates were used relative to bregma: SNc: (AP -2.3, ML ±1, DV -4.7), dorsal striatum: (AP +1.2, ML +1.8, DV -3.1). For more details, see dx.doi.org/10.17504/protocols.io.kqdg3x9j1g25/v1.

### Slice preparation

For *ex vivo* slice experiments, mice were anesthetized with isoflurane and then perfused with ice-cold cutting solution containing (in mM): 75 NaCl, 2.5 KCl, 6 MgCl_2_, 0.1 CaCl_2_, 1.2 NaH_2_PO_4_, 25 NaHCO_3_, 2.5 D-glucose, 50 sucrose and bubbled with 95% O_2_ and 5% CO_2_. Coronal slices (240 µm) containing the dorsal striatum and/or horizontal slices (240 µm) containing the SNc were cut in the same cutting solution. Slices were then incubated for 1 h at 32° C in artificial cerebrospinal fluid (aCSF) containing (in mM): 126 NaCl, 2.5 KCl, 1.2 MgCl_2_, 2 CaCl_2_, 1.2 NaH_2_PO_4_, 21.4 NaHCO_3_, 11.1 D-glucose and 10 µM MK-801 and bubbled with 95% O_2_ and 5% CO_2_. After incubation, slices were transferred into a recording chamber and constantly perfused with aCSF (33 ± 2 C) at a rate of 2 mL/min. Neurons were visualized using a BX51WI microscope (Olympus) with an infrared LED (Thorlabs). Fluorescent neurons were visualized with a custom-made green LED. Notably all mice that were used for ex vivo FSCV experiments were sacrificed following assessment of motor behavior in the open field. For more details, see dx.doi.org/10.17504/protocols.io.4r3l2omjpv1y/v1.

### Electrophysiology

All recordings were performed using an Axopatch 200B amplifier (Molecular Devices). Patch pipettes (2.5-4.5 MΩ) were made using a pipette puller (Narishige, PC-10). For experiments measuring D2- and GABA_A_-IPSCs simultaneously, pipettes for whole-cell recordings from MSNs contained (in mM): 67.5 D-gluconic acid (K), 67.5 KCl, 10 HEPES (K), 0.1 CaCl_2_, 2 MgCl_2_, 10 BAPTA, 0.1 mg/mL GTP, 1 mg/mL ATP, and 1.5 mg/mL phosphocreatine (pH 7.3, 280 mOsm). For experiments on ChIs, pipette solutions contained (in mM): 135 CsCl, 10 HEPES (K), 0.1 CaCl_2_, 2 MgCl_2_, 0.1 EGTA, 0.1 mg/mL GTP, 1 mg/mL ATP, and 1.5 mg/mL phosphocreatine (pH 7.35, 275 mOsm). For all other experiments, pipette solutions contained (in mM): 115 D-gluconic acid (K), 20 NaCl, 1.5 MgCl2, 10 HEPES(K), 10 BAPTA-tetrapotassium, 0.1 mg/mL GTP, 1 mg/mL ATP, and 1.5 mg/mL phosphocreatine (pH 7.3, 280 mOsm). All ChIs and MSNs were recorded in the dorsolateral corner of the dorsal striatum. Unless otherwise noted, aCSF contained CGP 55845 (300 nM), scopolamine (300 nM), dihydro-β-erythroidine (10 µM), and SKF 83566 (3 µM) and PTX (100 µM) for measuring AMPAR-EPSCs or DNQX (10 µM) for measuring GABA_A_A-IPSCs. Recordings of SNc dopamine neurons contained all the above blockers except for the GABA_B_R-IPSC experiments which included sulpiride (500 nM) in lieu of CGP 55845. Recordings of ChIs neurons contained all the above blockers including 500 nM sulpiride and 300 nM CGP 55845. Recordings were acquired with Axograph X (Axograph Scientific) at 10 kHz and filtered to 2 kHz. For whole-cell voltage-clamp recordings, cells were held at a voltage of -60 mV. No series resistance compensation was used, and a cell was discarded if its series resistance exceeded 15 MΩ. To activate ChR2-expressing dopamine neurons and their axons, 470 nm blue light (1 ms or 5 ms duration) was used to produce wide-field illumination. Prior to optical stimulation cells were held for a minimum of 4-5 minutes to allow for sufficient dialysis of the patched neuron. All recordings from GIRK2+ MSNs were performed in lateral regions of DStr showing robust tdTomato reporter fluorescence, which limits the variability of D2-receptor mediated GIRK outward currents between cells and among animals (Gong et al., 2021). For cholinergic interneuron recordings, slow mGluR1-mediated currents occasionally contained fast-inward currents. ChI recordings containing these fast-inward currents were excluded to more accurately determine current amplitudes. Expression of viral proteins were verified by post hoc histology on recorded slices. For more details, see dx.doi.org/10.17504/protocols.io.j8nlkooowv5r/v1.

### Fast scan cyclic voltammetry

Carbon fiber electrodes were encased with a glass pipette with an exposed diameter of 7 mm and length of 50 – 100 mm. The tip of fiber was placed in the DStr 30 – 70 mm below the surface of the slice. While holding the carbon fiber at 0.4 V, triangular waveforms (0.4 to 1.3 V versus Ag/AgCl at 400 V/s) were applied to the fiber at 10 Hz. Background subtracted cyclic voltammogram currents were obtained by subtracting the average of 10 voltammograms obtained prior to stimulation from each voltammogram obtained after stimulation. Peak [DA]_o_ was determined from the peak oxidation potential. The carbon fiber was calibrated to known concentrations of dopamine after experiments. For all voltammetry experiments, aCSF was used and dopamine release was evoked from DAT-Cre^(+/-)^, Ai32^(+/-)^ using a single 1 ms 470 nm blue light pulse. Expression of viral proteins were verified by post hoc histology on recorded slices. For more details, see dx.doi.org/10.17504/protocols.io.eq2lyj7krlx9/v1.

### Open field behavior

Locomotion was assessed using the open field test. For this assay, each mouse was acclimated in the behavior room for 30 minutes and then gently placed into a square arena (50 cm x 50 cm). Video recorded mouse movement for 30 min using an overhead camera. Tracking and post hoc analysis of total distance and velocity were conducted with custom MATLAB code. Expression of viral proteins were verified by post hoc histology on recorded slices. For more details, see dx.doi.org/10.17504/protocols.io.q26g7yo4kgwz/v1.

### Immunohistochemistry and fluorescence imaging

Mice were anesthetized using isoflurane and perfused transcardially with cold phosphate-buffered saline (PBS) followed by cold 4% paraformaldehyde in PBS (pH 7.4). Brains were post-fixed in 4% paraformaldehyde at 4°C for additional 24 hours, equilibrated in 30% sucrose solution for 2 days, and rapidly frozen in embedding freezing medium (Thermo Fisher Scientific). DSt and/or midbrain coronal slices of 50 μm in thickness were obtained using a Leica CM1950 cryostat (Leica Microsystems).

For total α-synuclein histology, sections were washed in PBS and incubated for 30 minutes at 80°C in 10 mM citrate buffer (antigen retrieval step). Slices were again washed in PBS prior to being blocked in 5% normal goat serum in PBS-T (0.3% Triton X-100) overnight at 4°C. Slices were then incubated with mouse anti-a-synuclein primary antibody (1:500; BD 610786) and rabbit anti-human a-synuclein antibody [MJFR1] primary antibody (1:500; Abcam ab138501) for 3 days at 4°C. After PBS washes, slices were incubated with goat anti-mouse Alexa Fluor 647 secondary antibody (1:500; Invitrogen Cat # A-21235) and goat anti-rabbit Alexa Fluor 488 secondary antibody (1:500; Invitrogen Cat # A-11008) overnight at 4°C and washed afterward with PBS. Sections were then mounted on slides for fluorescence imaging.

For TH immunohistochemistry, blocked in 5% normal goat serum in PBS-T for 2+ hour at room temperature (RT). Slices were then washed in PBS and incubated with rabbit anti-TH primary antibody (1:500; Millipore AB152) and rat α-synuclein (human) monoclonal antibody (15G7, 1:500 Enzo ALX-804-258) overnight at 4°C. After PBS washes, slices were incubated with goat anti-rabbit Alexa Fluor 488 secondary antibody (1:500; Invitrogen Cat # A-21245) and goat anti-rat Fluor 647 secondary antibody (1:500; Invitrogen Cat # A-11006) for 2+ hours at RT or overnight at at 4°C and washed afterward with PBS. Sections were then mounted on slides for fluorescence imaging.

For posthoc validation of viral expression, 240 μm coronal and horizontal sections were washed in cold PBS and then post fixed in cold 4% paraformaldehyde in PBS (pH 7.4) for a minimum of 24 h. Slices were then kept in in cold PBS at 4°C and blocked in 5% normal goat serum in PBS-T for 2+ hour at RT. Slices were then washed in PBS and incubated with rabbit anti-Alpha-synuclein antibody [MJFR1] primary antibody (1:500; Abcam ab138501) overnight at 4°C. After PBS washes, slices were incubated with goat anti-rabbit Alexa Fluor 647 secondary antibody (1:500; Invitrogen Cat # A-21245) for 1 hour at RT and washed afterward with PBS. Sections were then mounted on slice for fluorescence imaging. Only mice that showed mCherry or human α-synuclein within the striatum and/or midbrain of mice were included for analysis.

Following immunostaining, sections were finally mounted on slides with an anti-fade mounting media. For visualization of fluorescence reporters, sections were mounted without additional immunostaining. Fluorescent images were acquired using a slide scanner microscope (VS120, Olympus) and processed in Fiji (ImageJ, RRID:SCR_002285). For histological validation of total synuclein and TH in the α-synuclein overexpression model, human α-synuclein hemispheres were normalized against contralateral control/mCh-expressing hemispheres. For more details, see dx.doi.org/10.17504/protocols.io.14egn7yyzv5d/v1.

### Chemicals

Picrotoxin (1128), (+)-MK-801 maleate (0924), DNQX (0189), DHβE hydrobromide (2349), SKF 83566 hydrobromide (0925), (S)-(−)-sulpiride (0895), and scopolamine hydrobromide (1414) were obtained from Tocris Bioscience. EGTA (E4378) was from Sigma-Aldrich. CGP55845 hydrochloride (HB0960) was purchased from HelloBio and BAPTA tetra potassium salt was obtained from Invitrogen (B1204).

### Quantification and Statistical Analysis

All data are shown as mean ± SEM with individual points representing average values from individual animals. Statistical tests used for comparisons were parametric unpaired t test, non-parametric Mann-Whitney test, and Two-way ANOVA with Sidak post hoc tests where appropriate. Heteroscedasticity and normality assumptions of the data were tested to determine appropriate statistical tests for a given data set. The statistical significance was considered as p < 0.05 (*), p < 0.01 (**), p < 0.001 (***) and p < 0.001 (****). ‘N’ signifies number of animals and ‘n’ signifies number of cells, which can be found in the figure legends.

## Results

### Overexpression of human α-synuclein in midbrain dopamine neurons increases total synuclein levels and results in the onset of gross motor deficits

To examine the effects of wild-type human α-synuclein overexpression, we previously developed a cell-type specific adeno-associated viral strategy to express human α-synuclein together with mCherry (α-synuclein), or mCherry alone (mCherry) in a Cre-dependent manner (Barcomb et al., 2025). To further validate this model, we began by injecting AAV.DIO.human-α-synuclein.IRES.mCherry (AAV.hαSyn) into one hemisphere of the SNc of adult DAT-IRES-Cre^(+/-)^ mice, while injecting control AAV.DIO.IRES.mCherry (AAV.mCh) into the contralateral hemisphere (Figure 1A). As expected, this led to robust expression of human α-synuclein in the striatum of the AAV.α-synuclein injected hemisphere, both 4 and 8 weeks after viral transduction (Figure 1B). To determine the extent of overexpression of total α-synuclein, we next used a pan human and mouse α-synuclein antibody (syn-1) and quantified total α-synuclein levels in α-synuclein overexpressing hemispheres compared to mCherry controls. Consistent with our previous work (Barcomb et al., 2025), total α-synuclein fluorescence intensity, was increased approximately 1-fold in the dorsal striatum of hemispheres overexpressing α-synuclein compared to contralateral mCherry control hemispheres after both 4 and 8 weeks of expression (Figure 1B & 1C). Quantifying striatal TH expression, we found about a ∼30% decrease in TH levels in α-synuclein overexpressing hemispheres (Figure 1D & 1E). To further examine the extent of α-synuclein overexpression, we also quantified levels at the site of AAV injection within the SNc, finding approximately a 2-fold increase (Figure 1F & 1G). Despite the higher extent of α-synuclein overexpression, TH levels were similarly reduced in the SNc as in the striatum. As we previously found that α-synuclein overexpression with this viral approach does not lead to a loss of dopaminergic neurons within the SNc (Barcomb et al., 2025), the lowered TH levels in both the SNc and axon terminals likely results from a downregulation of TH expression, rather than a loss of dopamine neurons or axons.

**Figure 1:**
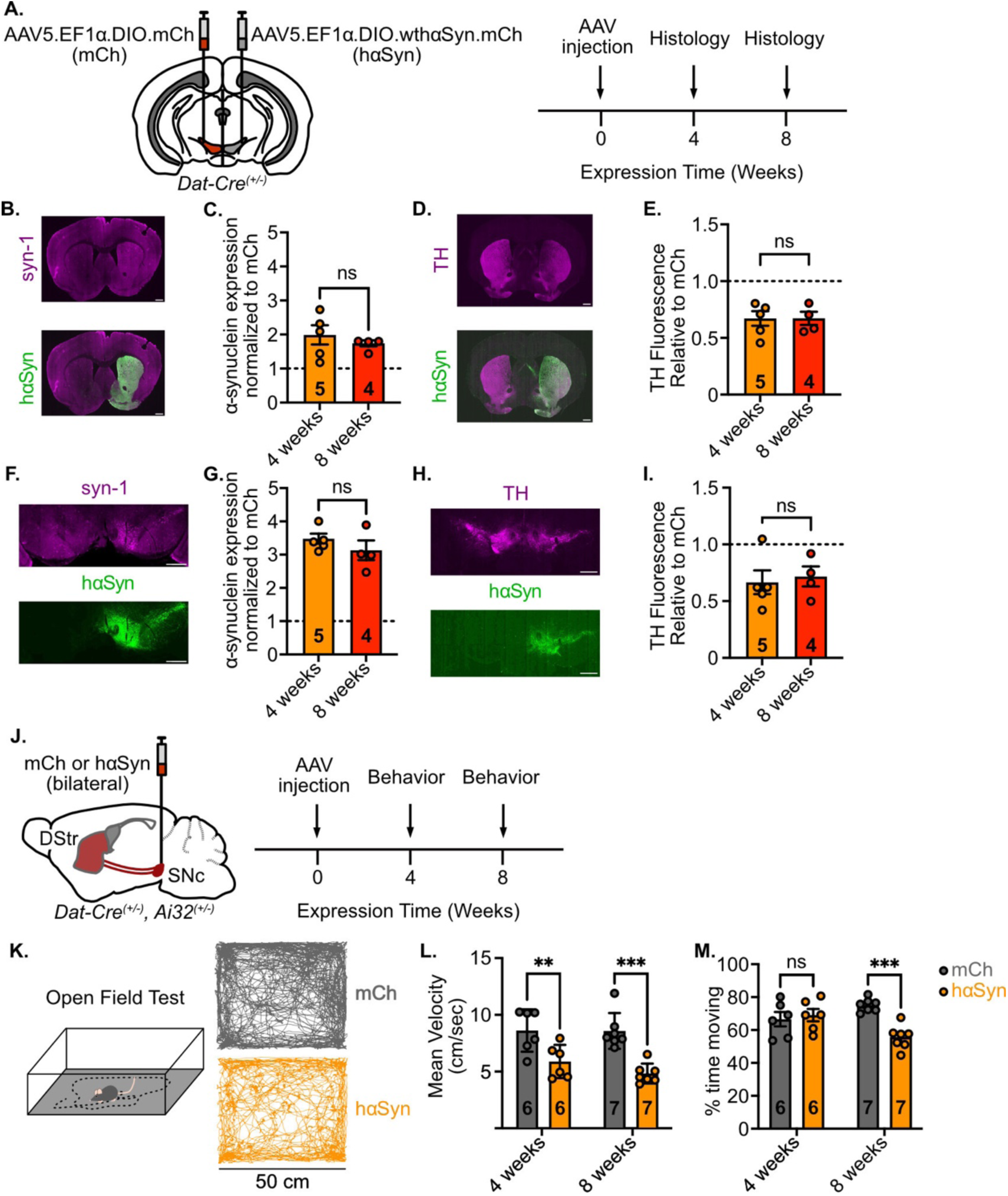
Histological and behavioral validation of the α-synuclein overexpression model. (A.) AAV-DIO-mCh and AAV-DIO-hαSyn-mCh were stereotaxically injected into contralateral SNcs of DAT-Cre^(+/-)^ mice and then sacrificed either 4 or 8 weeks after injection. (B.) Representative images of the dorsal striatum stained for total α-synuclein (Syn1, recognizes both human and mouse α-synuclein) and human α-synuclein. Scale bar: 500 μm. (C.) Quantification of total α-synuclein levels within the dorsal striatum of hαSyn-expressing hemispheres relative to the control/mCh injected hemispheres (Unpaired t-test). (D.) Representative images of the dorsal striatum stained for TH and hαSyn. Scale bar: 500 μm. (E.) Quantification of TH levels within the dorsal striatum of hαSyn-expressing hemispheres relative to the control/mCh injected hemispheres (Unpaired t-test). (F.) Representative images of the midbrain stained for total α-synuclein and hαSyn. Scale bar: 500 μm. (G.) Quantification of total α-synuclein levels within the midbrain of hαSyn-expressing hemispheres relative to the control/mCh injected hemispheres (Unpaired t-test). (H.) Representative images of the midbrain stained for TH and hαSyn. Scale bar: 500 μm. (I.) Quantification of TH levels within the midbrain of hαSyn-expressing hemispheres relative to the control/mCh injected hemispheres (Unpaired t-test). (J.) AAV-DIO-mCh or AAV-DIO-hαSyn-mCh were bilaterally injected into the SNcs of DAT-Cre^(+/-)^; Ai32^(+/-)^ mice and run on the open field either 4 or 8 weeks after injection. (K.) Representative traces showing mouse movement in the open field arena. (L.) hαSyn expression causes reductions in open field velocity after both 4 and 8 weeks of viral expression. (M.) α-synuclein expression causes reductions in open field velocity after 8 weeks, but not 4 weeks. (Two-way ANOVA with Sidak post hoc test)

Next, we examined the effect of α-synuclein overexpression on gross motor function. As subsequent physiology experiments required the ability to selectively stimulate dopamine terminals, we crossed DAT-IRES-Cre mice with Ai32 mice (DAT-Cre^(+/-)^ ; Ai32^(+/-)^) to express channelrhodopsin (ChR2) in dopamine cells and used this line for behavioral characterization. Mice were bilaterally injected into the SNc with AAV encoding either α-synuclein or mCherry control (Figure 1J). Overexpression of α-synuclein significantly decreased mean velocity of mice in the open field after 4 or 8 weeks compared to control mice (Figure 1K-M). Interestingly, percentage time moving was reduced in α-synuclein overexpressing mice after 8 weeks, but not 4 weeks of viral expression, which may explain why the velocity is more reduced at 8 weeks than at 4 weeks. This suggests that there is a progressive nature to motor deficits in our model with greater reductions in gross ambulation at later timepoints.

### α-synuclein overexpression reduces striatal bulk dopamine, but not D2R transmission within medium spiny neurons in the dorsolateral striatum

To determine the effect of α-synuclein overexpression on nigrostriatal dopamine transmission, we first used fast scan cyclic voltammetry (FSCV) in ex vivo brain slices to measure optogenetically evoked dopamine release in the striatum of DAT-Cre^(+/-)^ ; Ai32^(+/-)^ mice expressing either α-synuclein or mCherry. Dopamine release in the dorsolateral striatum (DLS) evoked by optical stimulation (1ms; 470 nm) was reduced in human α-synuclein expressing slices compared to mCherry controls at both 4- and 8-week timepoints (Figure 2B & 2C). As we previously found that α-synuclein overexpression does not alter striatal dopamine tissue content (Barcomb et al., 2025), the decrease in dopamine release driven by α-synuclein is likely due to direct effect on release rather than changes in dopamine levels as a result of reduced TH expression.

**Figure 2:**
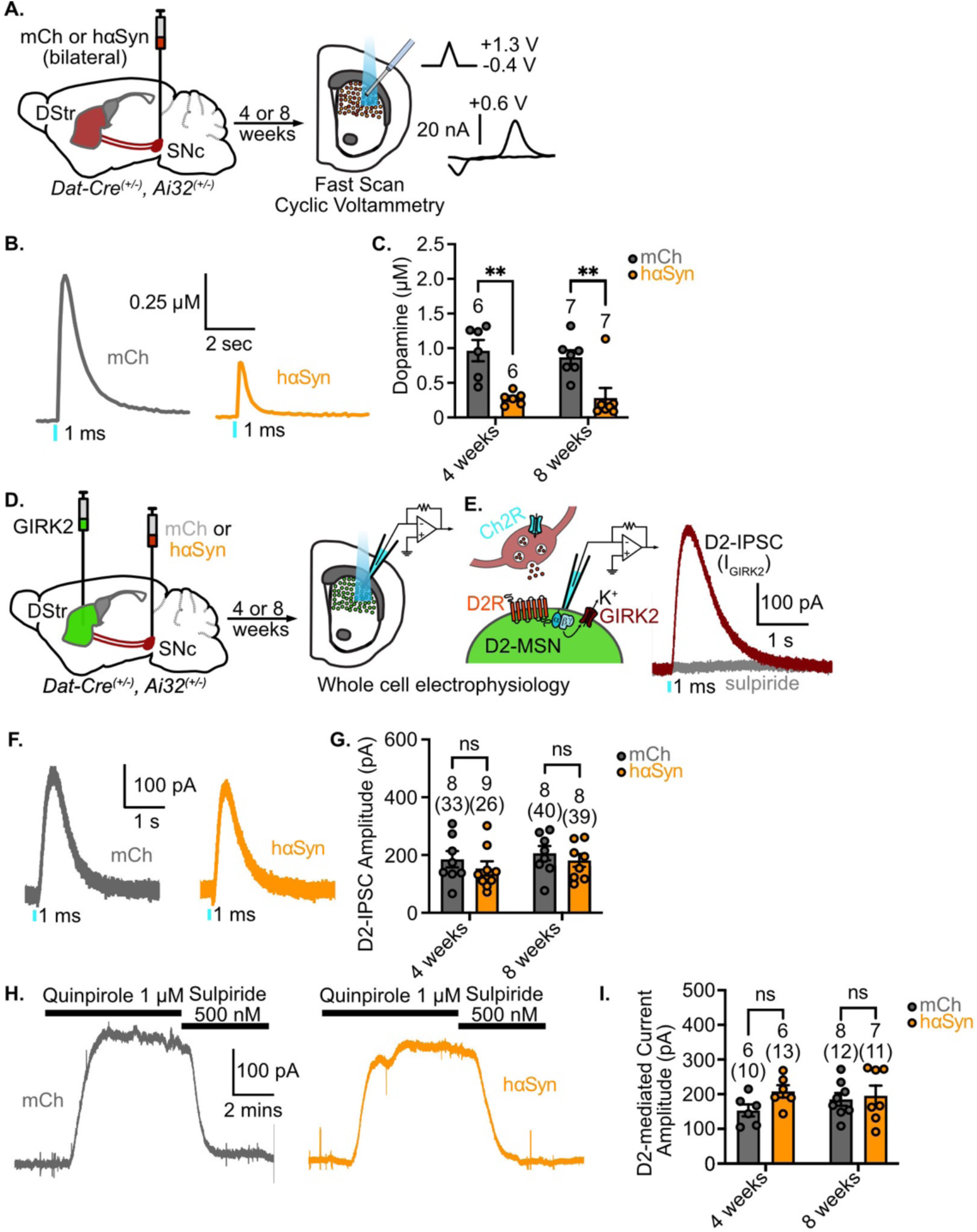
α-synuclein overexpression reduces striatal dopamine spillover, but not striatal dopamine transmission. (A.) AAV-DIO-mCh or AAV-DIO-hαSyn-mCh were bilaterally injected into the SNc of DAT-Cre^(+/-)^; Ai32^(+/-)^ mice and sacrificed for ex vivo voltammetry after 4 or 8 weeks of expression. (B.) Representative FSCV traces measuring the [DA]_o_ from a striatal brain slice in response to a single 1 ms optogenetic stimulation in either mCh or hαSyn injected mice. (C.) Quantification of optically-evoked changes in [DA]_o_ from mCh or hαSyn-expressing mice after 4 or 8 weeks of viral expression. (D.) AAV-DIO-mCh or AAV-DIO-hαSyn-mCh were injected into the SNcs and AAV-tdTomato-GIRK2 was injected into the dorsal striatum of DAT-Cre^(+/-)^; Ai32^(+/-)^ mice and sacrificed for ex vivo whole cell physiology after 4 or 8 weeks of expression. (E.) Representative traces of D2R-mediated GIRK currents from GIRK2+ D2-MSNs in response to a single 1 ms optogenetic stimulation. (F.) Representative D2-IPSCs in response to a single 1 ms optogenetic stimulation in either mCh or hαSyn injected mice. (G.) Quantification of optically-evoked D2-IPSCs from D2-MSNs in mCh or hαSyn expressing mice after 4 or 8 weeks of expression. (H.) Representative traces from GIRK2+ D2-MSNs shows that bath application of the quinpirole (1 μM) evokes an outward current that is reversed by sulpiride in either mCh or hαSyn injected mice. (I.) Quantification of quinpirole-evoked D2R-mediated currents from GIRK2+ D2-MSNs in mCh or hαSyn expressing mice after 4 or 8 weeks of expression (Two-way ANOVA with Sidak post hoc test).

While FSCV can be used to measure dopamine release, it does not measure direct dopamine signaling to post-synaptic striatal dopamine receptors. To examine synaptic dopamine transmission in the DLS, we next overexpressed G-protein coupled inwardly rectifying K^+^ (GIRK2) channels in striatal neurons, which efficiently couple to D2Rs in D2-receptor expressing MSNs and provide a robust electrophysiological readout of synaptic D2R activation (Marcott et al., 2014, 2018; Gong et al., 2021; Yee et al., 2025). As overexpressed GIRK channels do not couple to D1Rs (Marcott et al., 2014, 2018; Gong et al., 2021), the presence of D2R-mediated inhibitory postsynaptic currents (D2-IPSCs) provides a reliable readout of dopamine transmission by D2Rs in D2-MSNs, which are fully blocked following application of the D2R antagonist, sulpiride (500 nM) (Marcott et al., 2014; Gong et al., 2021) (Figure 2E). Using a cocktail of glutamate, GABA and acetylcholine receptor antagonists to pharmacologically isolate D2R signaling (Marcott et al., 2014, 2018; Gong et al., 2021), optogenetic stimulation reliably evoked D2-IPSCs in both human α-synuclein- and control mCherry-expressing slices (Figure 2F). While we saw a significant deficit in bulk dopamine release following α-synuclein overexpression, there was no difference in the amplitude of D2-IPSCs between α-synuclein overexpressing and mCherry control groups (Figure 2F & 2G). To test whether differential effects of α-synuclein overexpression on dopamine spillover (measured with FSCV) and receptor activation (D2-IPSCs) was the result of altered postsynaptic D2R signaling in MSNs, we measured whole-cell currents evoked by D2R agonist, quinpirole (1 µM) (Figure 2H & 2I). However, as the peak amplitude of quinpirole-evoked currents was similar between human α-synuclein- and mCherry-expressing slices (Figure 2H & 2I), this results suggests α-synuclein overexpression did not alter the efficacy of D2R signaling. Together, these results demonstrate that α-synuclein overexpression selectively impaired bulk dopamine release as detected by FSCV, with little effect on D2R activation in striatal D2-MSNs.

### GABA corelease from dopamine terminals in the DLS are not affected by α-synuclein expression

We next examined the effects of overexpressing α-synuclein on the co-transmission of GABA that is co-released from dopaminergic terminals in the DLS (Tritsch et al., 2012; Kim et al., 2015; Zych and Ford, 2022). To accomplish this, we pharmacologically isolated GABA_A_ receptors, and again recorded from MSNs in the DLS of DAT-Cre^(+/-)^ ; Ai32^(+/-)^ mice expressing either human α-synuclein or mCherry (Figure 3A). Optogenetic stimulation of dopamine terminals (1 ms, 470 nm) reliably evoked GABA_A_R-mediated inhibitory postsynaptic currents (GABA_A_R-IPSCs), which were abolished following application of the GABA_A_R antagonist, picrotoxin (Figure 3B). In order to distinguish D1-MSNs from D2-MSNs we took advantage of the fact that D2Rs but not D1Rs efficiently couple to GIRK channels (Marcott et al., 2014, 2018; Gong et al., 2021) by virally expressing GIRK channels in striatal neurons and used the subsequent presence or absence of D2-IPSCs to identify cells as either D2-MSNs or D1-MSNs respectively. We found no significant difference in the amplitude of GABA_A_R-IPSCs in D2-MSNs in slices from α-synuclein overexpressing mice compared to mCherry controls at both 4- and 8-week timepoints (Figure 3D & 3E). Likewise, GABA_A_R-IPSCs recorded from D1-MSNs were also similar between human α-synuclein and mCherry control groups (Figure 3G & 3H). Thus, GABA corelease within the DLS appears to be unaffected by α-synuclein overexpression.

**Figure 3:**
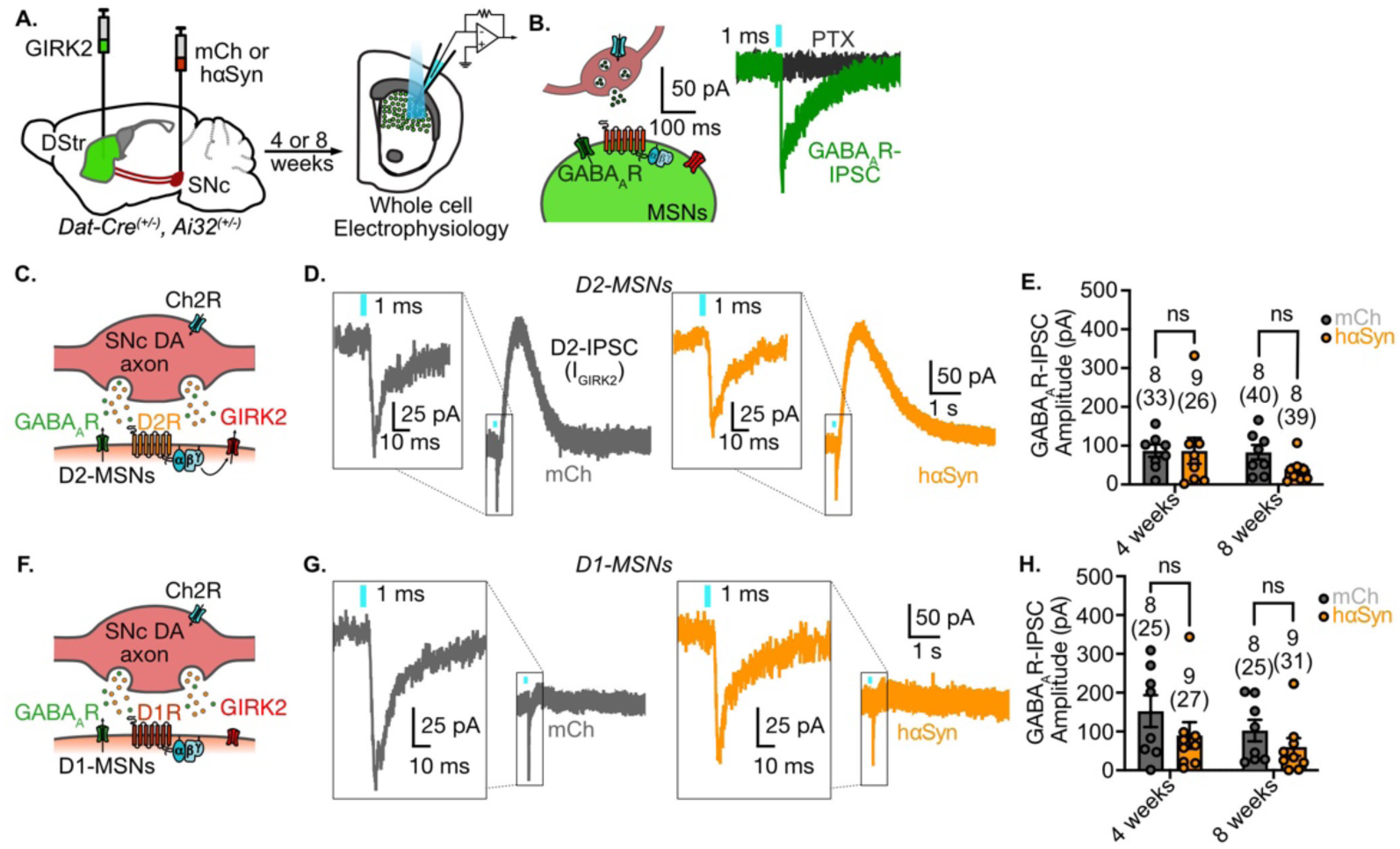
α-synuclein overexpression does not alter GABA corelease from dopamine terminals in the DLS. (A.) AAV-DIO-mCh or AAV-DIO-hαSyn-mCh were injected into the SNc and AAV-tdTomato-GIRK2 was injected into the dorsal striatum of DAT-Cre^(+/-)^; Ai32^(+/-)^ mice and sacrificed for ex vivo whole cell physiology after 4 or 8 weeks of expression. (B.) Optical stimulation of SNc dopamine terminals within the DLS evokes GABA release that can be measured by whole cell patch clamp electrophysiology. (C.) Optical stimulation of SNc dopamine terminals within the DLS results in GABA_A_R-IPSCs in D2-MSNs. (D.) Optically evoked GABA_A_R-IPSCs and D2-IPSCs in D2-MSNs from mCh/control and hαSyn-expressing mice. (E.) Quantification of optically-evoked GABA_A_R-IPSCs from D2-MSNs in mCh or hαSyn expressing mice after 4 or 8 weeks of expression. (F.) Optical stimulation of SNc dopamine terminals within the DLS results in GABA_A_R-IPSCs in D1-MSNs. (G.) Optically evoked GABA_A_R-IPSCs in D1-MSNs from mCh/control and hαSyn-expressing mice. (H.) Quantification of optically-evoked GABA_A_R-IPSCs from D1-MSNs in mCh or hαSyn expressing mice after 4 or 8 weeks of expression. (Two-way ANOVA with Sidak post hoc test)

### α-synuclein overexpression reduces glutamate corelease from dopamine terminals in the DLS

Next, we proceeded to examine the effects of overexpressing α-synuclein on glutamate co-transmission, first examining glutamate co-release onto MSNs. In the presence of antagonists to pharmacologically isolate AMPA receptors (AMPARs), optogenetic stimulation of dopamine terminals evoked AMPAR excitatory postsynaptic currents (AMPA-EPSCs) in DLS MSNs, which were blocked by the AMPAR antagonist, DNQX (Figure 4A-4B). Similar to the approach for measuring GABA co-transmission, we expressed GIRK channels in MSNs and again used the presence or absence of D2-IPSCs to identify neurons as either D2-MSNs or D1-MSNs. In contrast to GABA corelease, the amplitude of AMPAR-EPSCs was significantly reduced in slices expressing human α-synuclein compared to slices from mCherry control animals after 4 weeks and 8 weeks of expression in both D2-MSNs and D1-MSNs (Figure 4C – 4H). Thus, overexpression of α-synuclein appears to have a selective effect on glutamate corelease from dopamine terminals in the DLS.

**Figure 4:**
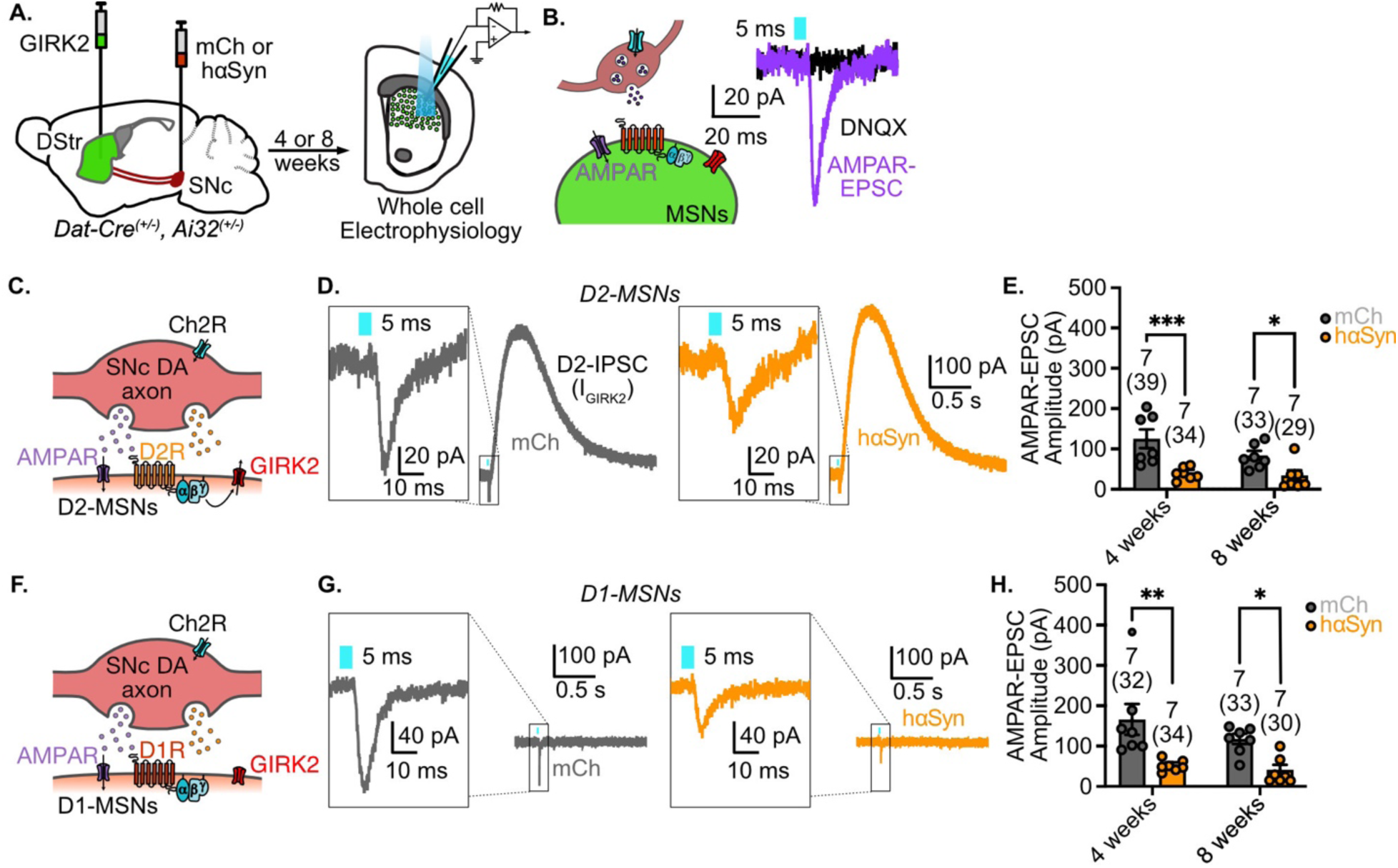
α-synuclein overexpression reduces glutamate corelease from dopamine terminals onto MSNs in the DLS. (A.) AAV-DIO-mCh or AAV-DIO-hαSyn-mCh were injected into the SNc and AAV-tdTomato-GIRK2 was injected into the dorsal striatum of DAT-Cre^(+/-)^; Ai32^(+/-)^ mice and sacrificed for ex vivo whole cell physiology after 4 or 8 weeks of expression. (B.) Optical stimulation (5 ms) of SNc dopamine terminals within the DLS evokes glutamate release that can be measured by whole cell patch clamp electrophysiology. (C.) Optical stimulation of SNc dopamine terminals within the DLS results in AMPAR-EPSC in D2-MSNs. (D.) Optically evoked AMPAR-EPSCs and D2-IPSCs in D2-MSNs from mCh/control and hαSyn-expressing mice. (E.) Quantification of optically-evoked GABA_A_R-IPSCs from D2-MSNs in mCh or hαSyn expressing mice after 4 or 8 weeks of expression. (F.) Optical stimulation of SNc dopamine terminals within the DLS results in AMPAR-EPSCs in D1-MSNs. (G.) Optically evoked GABA_A_R-IPSCs in D1-MSNs from mCh/control and hαSyn-expressing mice. (H.) Quantification of optically-evoked GABA_A_R-IPSCs from D1-MSNs in mCh or hαSyn expressing mice after 4 or 8 weeks of expression. (Two-way ANOVA with Sidak post hoc test)

In addition to transmission to MSNs, SNc dopamine axons also co-release glutamate onto DLS cholinergic interneurons (ChIs) to mediate burst firing via metabotropic glutamate receptor (mGluR1) signaling (Straub et al., 2014; Cai and Ford, 2018; Chuhma et al., 2018). To determine whether the reduction in glutamate corelease that we see in striatal MSNs was also evident across different neurons, we recorded from DLS ChIs and isolated mGluR1-mediated currents evoked with a burst of optical stimulation (5 stimuli at 20 Hz, 2 ms) (Figure 5B). Slow mGluR1-mediated inward currents were reliably recorded in ChIs as has been seen previously (Cai and Ford, 2018; Chuhma et al., 2018) (Figure 5C). Similar to the effects of α-synuclein overexpression in MSNs, the amplitude of mGluR1 currents in ChIs was reduced in α-synuclein overexpressing mice compared to mCherry controls at both 4- and 8-week timepoints (Figure 5D). Taken together, these results indicate that glutamate corelease from dopamine terminals in the DLS exhibits a selective sensitivity to α-synuclein overexpression that is not evident in synaptic-like release of dopamine or GABA corelease.

**Figure 5:**
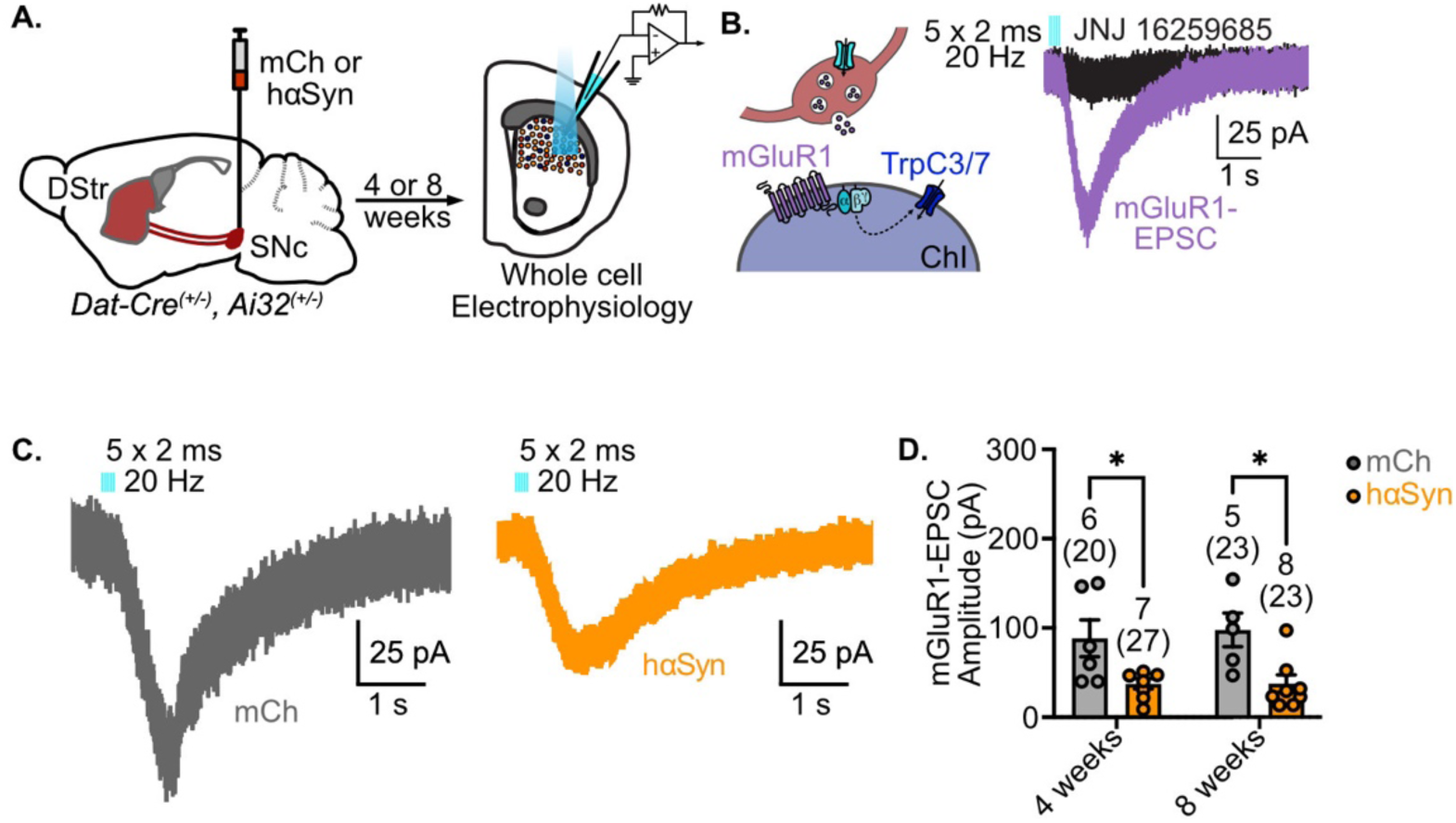
α-synuclein overexpression impairs glutamate corelease onto ChIs in the DLS. (A.) AAV-DIO-mCh or AAV-DIO-hαSyn-mCh were injected into the SNc of Dat-Cre^(+/-)^; Ai32^(+/-)^ mice and sacrificed for ex vivo whole cell physiology after 4 or 8 weeks of expression. (B.) Optical stimulation (5 x 2 ms at 20 Hz) of SNc dopamine terminals within the DLS evokes glutamate release onto ChIs can be measured by whole cell patch clamp electrophysiology. Optically-evoked mGluR1-mediated EPSCs are blocked by 20 μM JNJ 16259685. (C.) Representative traces of mGluR1-mediated EPSCs in mCh and hαSyn injected mice. (D.) Quantification of mGluR1-mediated EPSCs from DLS ChIs in mCh- or hαSyn-expressing mice after 4 or 8 weeks of expression. (Two-way ANOVA with Sidak post hoc test)

### Somatodendritic dopamine release within the SNc is abolished by α-synuclein overexpression

In addition releasing dopamine from axon terminals, SNc dopamine neurons also release dopamine from somatodendritic sites (Björklund and Lindvall, 1975; Geffen et al., 1976; Jaffe et al., 1998). The somatodendritic release of dopamine mediates pauses in SNc neuron firing as a result of D2R activation of GIRK channels and a resulting D2R-mediated inhibitory postsynaptic current (D2-IPSC) (Beckstead et al., 2004; Ford et al., 2010; Gantz et al., 2013; Hikima et al., 2021). As loss of somatodendritic dopamine release has been proposed to partially underlie motor deficits following dopamine neuron degeneration (Bergquist et al., 2003; González-Rodríguez et al., 2021), we next examined the effect of overexpressing α-synuclein on somatodendritic dopamine transmission.

SNc dopamine cells from DAT-Cre^(+/-)^ ; Ai32^(+/-)^ mice were identified based on morphology, the presence of pacemaker-like firing, the presence of inward hyperpolarization-activated (I_H_) currents and direct ChR2-mediated currents (Grace and Onn, 1989; Mercuri et al., 1995). We first examined the effects of α-synuclein overexpression on SNc neuron intrinsic properties and excitability. Consistent with past findings (Barcomb et al., 2025), overexpression of α-synuclein had minimal impact on SNc dopamine neurons spontaneous firing rate (Figure 6C), coefficient of variability (Figure 6D), action potential half width (Figure 6E) or I_H_ current amplitudes (Figure 6F & 6G). These recordings indicate that α-synuclein overexpression, using the current model, has little effect on the excitability of dopamine neurons themselves.

**Figure 6:**
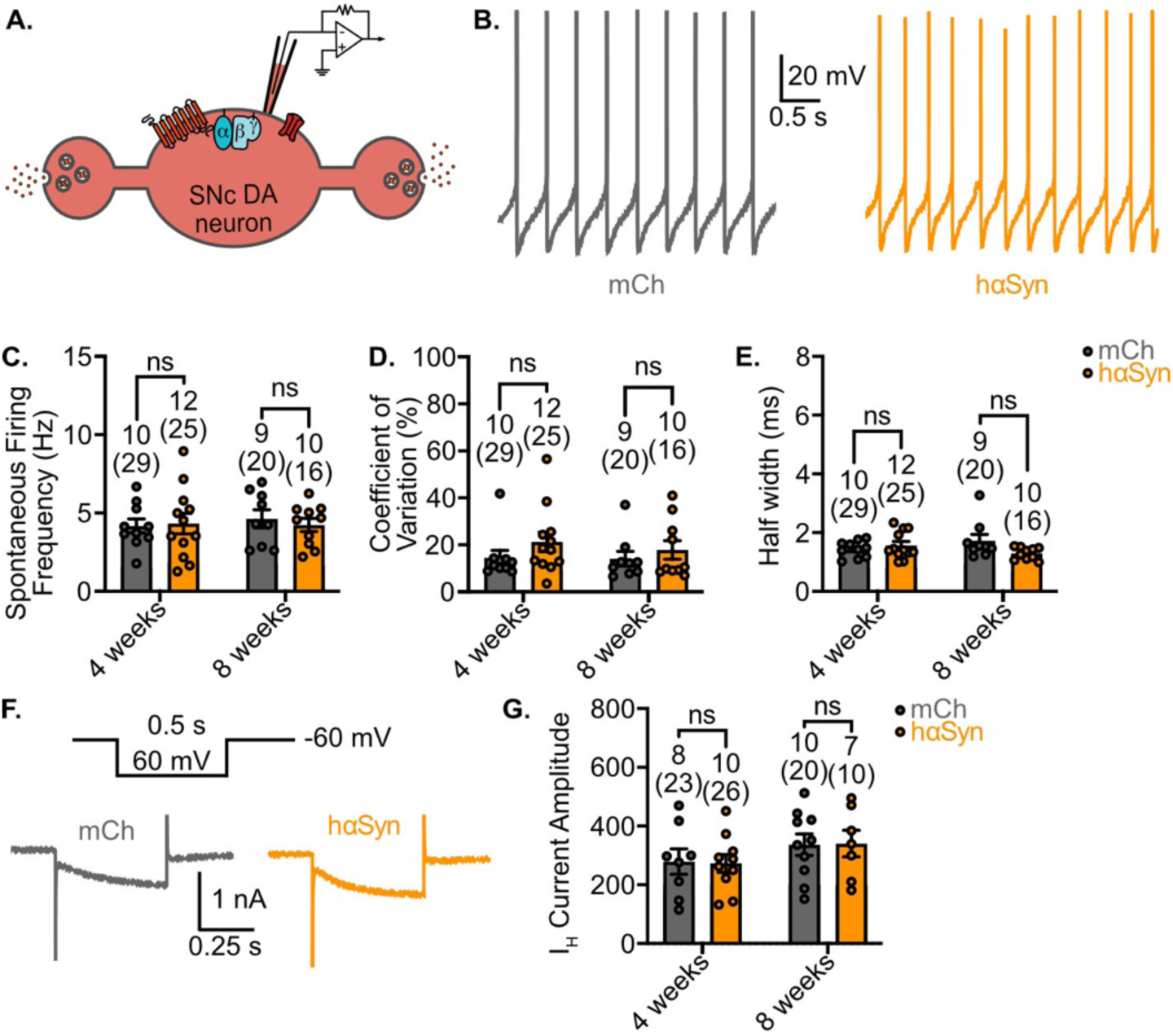
α-synuclein overexpression does not affect intrinsic properties of SNc DA neurons. (A.) AAV-DIO-mCh or AAV-DIO-hαSyn-mCh were injected into the SNc of Dat-Cre^(+/-)^; Ai32^(+/-)^ mice and sacrificed for ex vivo whole cell physiology after 4 or 8 weeks of expression. (B.) Representative traces of SNc DA neurons spontaneous firing in mCh and hαSyn injected mice. Quantification of spontaneous action potential firing frequency (C.), coefficient of variation (D.), and half width (E.) in mCh or hαSyn-expressing mice after 4 or 8 weeks of expression. (F.) Representative hyperpolarization-activated K+ currents (I_H_) evoked by a 60-mV hyperpolarizing step in mCh and hαSyn-expressing mice. (G.) Quantification of I_H_ evoked by a 60-mV hyperpolarizing step in cells from mCh or hαSyn injected mice after 4 or 8 weeks of expression. (Two-way ANOVA with Sidak post hoc test)

Next, we measured somatodendritic transmission onto SNc dopamine neurons evoked by trains of electrical stimuli (5x 30 µA, 0.5 ms) provided from a nearby monopolar stimulating electrode (Figure 7A & 7B). Electrical stimulation evoked somatodendritic D2-IPSCs in mCherry expressing control SNc neurons, which were blocked by sulpiride (Figure 7B). Strikingly, somatodendritic D2-IPSCs were completely absent in SNc neurons overexpressing α-synuclein (Figure 6C & 6D). This loss of somatodendritic transmission was evident after both 4 and 8 weeks of expression. To determine whether the loss of D2-IPSCs resulted from a presynaptic deficit of dopamine release or a postsynaptic alteration in D2R signaling, we exogenously activated D2Rs by bath application of D2R agonist quinpirole (1 μM). The amplitude of quinpirole-evoked currents was similar in human α-synuclein and mCherry control cells (Figure 7E & 7F), indicating that α-synuclein overexpression impacted presynaptic release, as opposed to D2R function. To confirm that α-synuclein overexpression selectively affected somatodendritic dopamine transmission, we also examined electrically evoked GABA_B_ receptor mediated IPSCs (GABA_B_-IPSCs; Figure 7G & 7H) as GABA_B_R receptors similarly couple to GIRK channels in SNc neurons to mediate a metabotropic IPSC (Lacey et al., 1988). In contrast to somatodendritic D2-IPSCs, GABA_B_-IPSCs were completely unaffected by α-synuclein overexpression (Figure 7H & 7I). Taken together, these findings suggest that in contrast to striatal dopamine transmission, α-synuclein overexpression selectively affects somatodendritic dopamine release, potentially by affecting unique release machinery that is utilized for somatodendritic dopamine release in this region (Witkovsky et al., 2009; Rice and Patel, 2015; Hikima et al., 2022).

**Figure 7:**
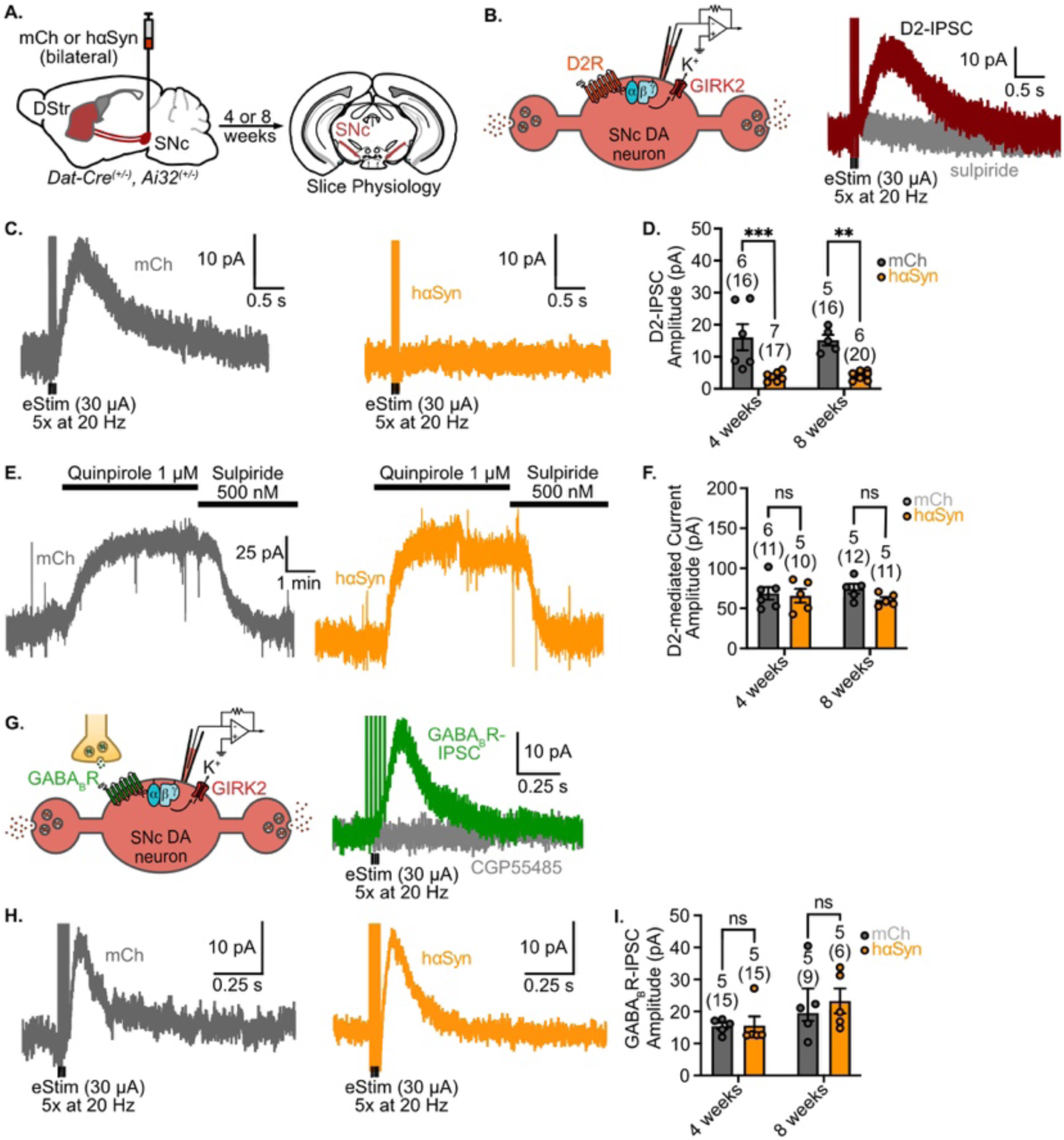
α-synuclein overexpression abolishes somatodendritic DA release within the SNc. (A.) AAV-DIO-mCh or AAV-DIO-hαSyn-mCh were injected into the SNc of DAT-Cre^(+/-)^; Ai32^(+/-)^ mice and sacrificed for ex vivo whole cell physiology after 4 or 8 weeks of expression. (B.) Electrical stimulation illicits dopamine release from somatodendritic compartments which can be measured using whole cell electrophysiology. Electrically-evoked somatodendritic D2-IPSCs are blocked by 500 nM sulpiride. (C.) Representative traces of electrically-evoked (5x30 μA at 20 Hz) somatodendritic D2-IPSCs in cells from mCh/control or hαSyn-expressing mice. (D.) Quantification of somatodendritic D2-IPSCs from SNc dopamine neurons in mCh or hαSyn expressing mice after 4 or 8 weeks of expression. (E.) Representative traces from SNc DA neurons shows that bath application of the quinpirole (1 μM) evokes an outward current that is reversed by sulpiride in both mCh or hαSyn injected mice. (F.) Quantification of quinpirole-evoked D2R-mediated currents from SNc DA neurons in mCh- or hαSyn-expressing mice after 4 or 8 weeks of expression. (G.) Electrical stimulation elicits dopamine release from local GABA terminals within the SNc, pharmacological isolation of GABA_B_Rs expressed by SNc DA neurons allows measurement of GABA_B_R currents in the whole cell configuration. (H.) Representative traces of electrically-evoked (5x30 μA at 20 Hz) GABA_B_R-IPSCs in SNc DA neurons from mCh/control or hαSyn-expressing mice. Electrically-evoked GABA_B_R-IPSCs are blocked by 300 nM CGP55845. (I.) Quantification of GABA_B_R-IPSCs from SNc dopamine neurons in mCh- or hαSyn-expressing mice after 4 or 8 weeks of expression. (Two-way ANOVA with Sidak post hoc test)

## Discussion

Here we examined the effects of α-synuclein overexpression on nigrostriatal transmission prior to dopaminergic cell loss. We found that cell-type specific overexpression of wild-type human α-synuclein in midbrain dopamine neurons selectively impaired glutamate co-release from axon terminals in the dorsal striatum, and completely abolished somatodendritic dopamine transmission within the SNc. In contrast, we found that GABA corelease from dopamine axons was unaffected by α-synuclein overexpression, as was the extent of D2R activation in striatal MSNs (see Figure 8). These findings indicate that different aspects of nigrostriatal transmission are selectively impacted by α-synuclein, which may contribute to motor dysfunction caused by α-synuclein accumulation in early stages of PD progression.

**Figure 8:**
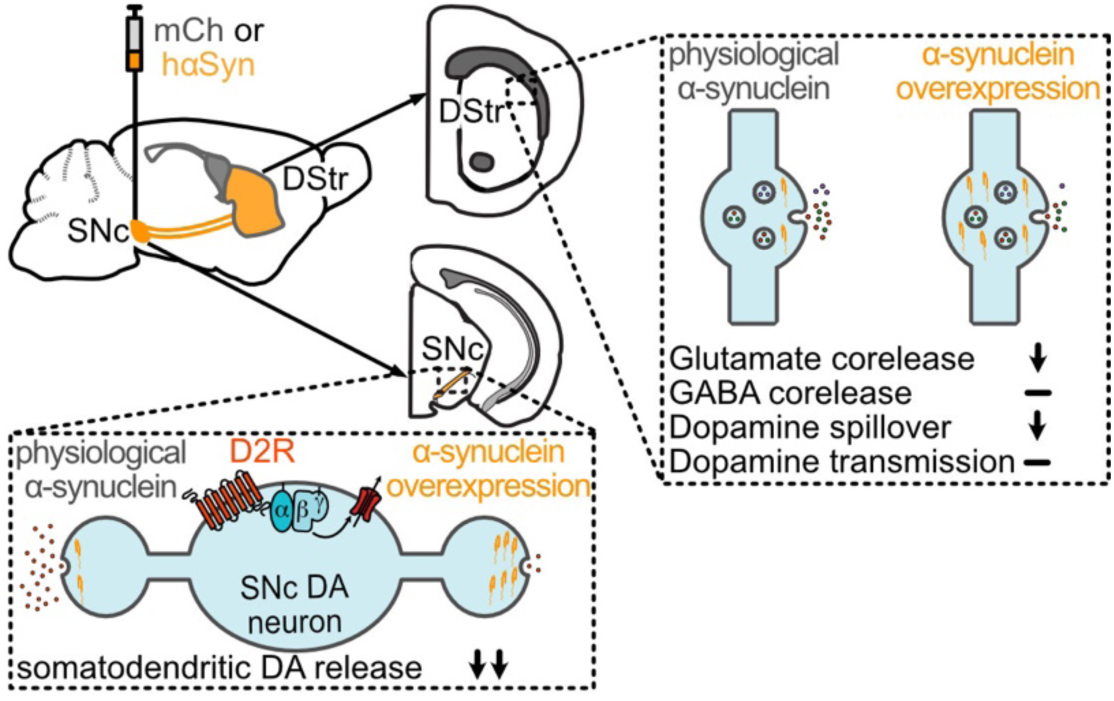
Summary figure showing the nigrostriatal release deficits associated with α-synuclein overexpression.

### Effects of α-synuclein overexpression on striatal dopamine release and D2R transmission

Consistent with past studies (Lundblad et al., 2012; Janezic et al., 2013; Barcomb et al., 2025), we found that α-synuclein overexpression reduced dopamine release in the dorsal striatum. Despite this reduction in dopamine release, D2R activation, when measured via GIRK currents in D2-MSNs, was not significantly different between human α-synuclein and mCherry control groups. While increased D2R expression has been reported in PD patients (Ryoo et al., 1998), we did not see changes in the amplitude of quinpirole-evoked currents in MSNs following α-synuclein overexpression suggesting that differences in receptors sensitivity did not occur. Instead, as FSCV and D2R activation differ in their sensitivity for dopamine, differences in measurements may partly account for this observation (Ford et al., 2009; Marcott et al., 2014; Yee et al., 2025). In addition, since the distance from the carbon fiber electrode to dopamine release sites is orders of magnitude greater than the distance between membrane-bound postsynaptic D2Rs and release sites, it is also possible that this more proximal dopamine release may be less affected by α-synuclein overexpression. Future work using membrane-expressed, genetically-encoded fluorescent sensors may shed light on this disparity. Finally, compensatory adaptations within the nigrostriatal system may also contribute, as such mechanisms are well established in PD and are thought to maintain dopaminergic signaling despite progressive neuron loss (Bezard et al., 2003). Whether changes in dopamine reuptake, autoreceptor sensitivity, or other factors help preserve local D2R transmission in this context remains to be determined.

### Effect of increased α-synuclein on GABA and glutamate co-release from SNc dopamine terminals

We found a differential effect of α-synuclein overexpression on glutamate and GABA co-release from SNc dopamine terminals in the DLS. Glutamate co-release onto both MSNs and cholinergic interneurons was robustly reduced in α-synuclein overexpressing animals, while GABA co-release onto D1- and D2-MSNs was unaffected. This is in contrast to previous observations that mutant α-synuclein (A53T) more strongly affected GABA co-transmission (Kim et al., 2023) but is consistent with reports that α-synuclein overexpression induces glutamatergic deficits that precede reductions in GABA transmission (Wu et al., 2010), suggesting that glutamatergic release machinery may be particularly sensitive to elevated α-synuclein levels.

The selective sensitivity of glutamate co-release to α-synuclein overexpression may reflect fundamental differences in the regulation of different pools of vesicles in dopamine terminals. Within SNc terminals in the dorsolateral striatum, glutamate is packaged into vesicles by VGluT2, whereas GABA is packaged by the vesicular monoamine transporter VMAT2 (Tritsch et al., 2012; Melani and Tritsch, 2022; Zych and Ford, 2022). Since VGluT2- and VMAT2-containing vesicles have been shown to largely segregate into distinct vesicular pools with different release properties (Silm et al., 2019), α-synuclein overexpression may preferentially disrupt VGluT2-mediated glutamate packaging or release, while leaving VMAT2-mediated GABA co-release and VMAT2-dependent dopamine transmission to postsynaptic D2Rs in MSNs comparatively intact. How α-synuclein accumulation selectively disrupts VGluT2-containing vesicle pools within dopamine terminals is an important question for future investigation.

### α-synuclein regulation of somatodendritic dopamine release within the SNc

Using this model, we found that α-synuclein overexpression completely abolished somatodendritic D2-IPSCs in SNc dopamine neurons after both 4 and 8 weeks of expression. To determine whether this loss reflected a presynaptic deficit in dopamine release or a postsynaptic change in D2R signaling, we applied the D2R agonist quinpirole directly and found that D2R-mediated outward currents were comparable between α-synuclein overexpressing and mCherry control neurons. Furthermore, GABA_B_-IPSCs evoked by local electrical stimulation were unaffected by α-synuclein overexpression, confirming that GPCR-to-GIRK coupling in SNc dopamine neurons is intact. Taken together, these data establish that α-synuclein overexpression selectively abolishes somatodendritic dopamine release without altering the downstream postsynaptic signaling machinery.

Notably, α-synuclein overexpression did not affect the intrinsic firing properties of SNc dopamine neurons, indicating that the loss of somatodendritic release reflects a direct disruption of the release process rather than a secondary consequence of altered excitability. The somatodendritic compartment utilizes release machinery that is distinct from that at axonal terminals (Kennedy and Ehlers, 2011; Rice and Patel, 2015; Ludwig et al., 2016; Robinson et al., 2019; Hikima et al., 2022; Lebowitz et al., 2023; Choi et al., 2026), and the complete loss of somatodendritic release we observe here suggests that this unique machinery is particularly vulnerable to wild-type α-synuclein accumulation. Interestingly, this pattern differs markedly from mitochondrial models of PD. In mice with dopamine neuron-specific Ndufs2 deficiency (MCI Park), axonal dopamine release is severely impaired at timepoints where somatodendritic release is relatively preserved (González-Rodríguez et al., 2021). This contrast suggests that α-synuclein accumulation and mitochondrial dysfunction impair dopamine release through mechanistically distinct pathways, with accumulated α-synuclein preferentially disrupting the somatodendritic release machinery.

Somatodendritic dopamine release activates D2 autoreceptors on SNc dopamine neurons, mediating inhibitory pauses in firing that regulate overall dopamine output (Beckstead et al., 2004; Ford et al., 2010; Gantz et al., 2013; Hikima et al., 2021). Beyond this local autoregulatory role, somatodendritic dopamine release has been proposed to contribute to gross motor control, and deficits in somatodendritic release have been linked to the emergence of locomotor impairments in progressive mitochondrial-based models of PD (Bergquist et al., 2003; González-Rodríguez et al., 2021; Muñoz et al., 2025). The complete loss of somatodendritic release in our model, in the absence of changes to intrinsic properties or postsynaptic signaling, may therefore represent a particularly relevant contributor to motor dysfunction arising from α-synuclein accumulation.

### Nigrostriatal determinants of PD-related motor phenotypes

In our α-synuclein overexpression model, deficits in bulk striatal dopamine release, glutamate co-release, and somatodendritic dopamine release all parallel the onset of gross motor impairment. While it is difficult to establish a direct causal link between any individual synaptic deficits and motor dysfunction, prior work provides useful context for interpreting the relative contributions of each. Although loss of glutamate co-release specifically onto cholinergic interneurons within the DLS has been shown to drive PD-related motor deficits (Cai et al., 2021), broader evidence suggests that glutamate co-release does not play a major role in gross motor function (Hnasko et al., 2010; Fortin et al., 2012; Nordenankar et al., 2015; Mingote et al., 2017). It therefore seems unlikely that reduced glutamate co-release alone drives the locomotor reductions observed here.

Our finding that reduced bulk striatal dopamine parallels locomotor deficits is consistent with results from a genetic synuclein overexpression model (SNCA-OVX), in which constitutive human α-synuclein expression reduces gross ambulation (Janezic et al., 2013). Notably, in that model, striatal dopamine spillover deficits precede the onset of motor impairment, suggesting that the initial reduction in dopamine release may not in itself be sufficient to drive locomotor dysfunction (Janezic et al., 2013). Whether this is also the case in our model is not clear as the earliest timepoint that we examined was at 4 weeks, at which point deficits in dopamine release and locomotion were apparent. The combined loss of striatal dopamine spillover and the complete abolition of somatodendritic dopamine release observed here may together contribute to the motor deficits in our model, though future studies examining the temporal emergence of each deficit relative to motor impairment will be needed to better resolve their individual contributions.

More broadly, our findings highlight the importance of studying α-synuclein accumulation in isolation from other pathogenic processes to understand its specific contributions to nigrostriatal dysfunction. Wild-type α-synuclein accumulation can drive motor dysfunction and selective synaptic deficits in the absence of dopaminergic cell loss and terminal degeneration, providing a window into the earliest functional consequences of PD-relevant pathology. Complementary models targeting mitochondria, the endo-lysosomal system, and α-synuclein will each be important for dissecting the distinct mechanisms that drive different aspects of PD progression. By establishing how each process contributes to specific synaptic and behavioral deficits, we can build a more complete picture of early PD pathology and identify which aspects of nigrostriatal dysfunction are most relevant as targets for intervention at the pre-degenerative stage of disease.

## Data and Materials Availability

All generated data is available in the main text, Supplemental Table S1 or at Zenodo repository (https://doi.org/10.5281/zenodo.20517296).

The data, code, protocols, and key lab materials used in this study are listed in a Key Resource Table alongside their persistent identifiers at Zenodo repository https://doi.org/10.5281/zenodo.20517296).

No code was generated for this study; all data cleaning, preprocessing, analysis, and visualization was performed using commercial software stated in Materials and Methods.

## Author contributions

NB, AGY and CPF designed experiments. RHE generated the human α-synuclein and control mCherry viruses. NB and AGY performed research experiments and analyzed the data. NB and CPF wrote the manuscript.

## Acknowledgments

This work was funded by NIH grants R01-NS138043 (CPF), as well as funded in part by Aligning Science Across Parkinson’s (ASAP-020529) (CPF & RHE) through the Michael J. Fox Foundation for Parkinson’s Research (MJFF). For the purpose of open access, the author has applied a CC BY public copyright license to all Author Accepted Manuscripts arising from this submission.

---

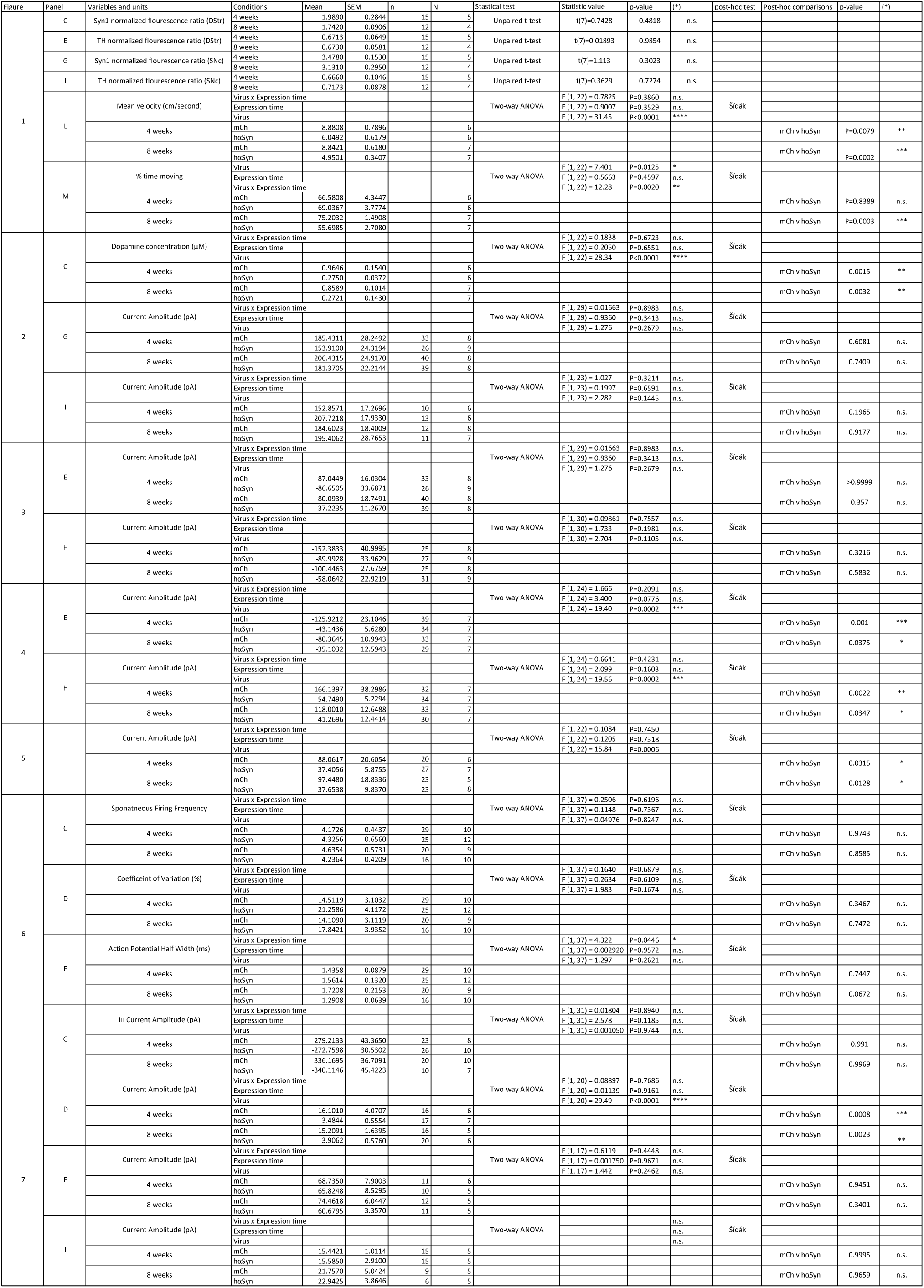

